# RNA Sequencing Reveals Divergent Transcriptome Changes in SBP2 and SECISBP2L Depleted Cell Lines

**DOI:** 10.1101/2025.07.02.662884

**Authors:** Jesse Donovan, Paul R. Copeland

## Abstract

Selenocysteine (Sec), the 21st amino acid, is co-translationally inserted at UGA codons via a specialized machinery requiring SECIS elements, Sec-tRNA^Sec, eEFSec, and SECIS-binding protein 2 (SBP2). While SBP2 is essential for Sec incorporation in vitro and in vivo, the function of its paralog, SECISBP2L, remains incompletely defined. In this study, we investigated the distinct roles of SBP2 and SECISBP2L in the human hepatocellular carcinoma cell line HepG2, which expresses a broad selenoproteome. Using CRISPR-Cas9 genome editing, we generated SBP2 and SECISBP2L edited cell lines. Consistent with previous findings, SBP2 targeting impaired selenoprotein mRNA and protein expression, whereas SECISBP2L targeting did not. However, transcriptomic profiling by RNA-seq revealed that SECISBP2L targeting induced differential expression of over 800 genes, with significant enrichment in pathways related to extracellular matrix organization and cell adhesion. In contrast, SBP2 targeting produced a distinct transcriptomic signature enriched for metabolic and ion transport processes. Notably, only limited overlap in differentially expressed genes was observed between the two knockout models. Mass spectrometry and immunoblot data indicated that CRISPR-targeted SECISBP2L cells produce a truncated protein via internal translation initiation, suggesting that observed gene expression changes may be attributable to loss of a portion of the SECISBP2L N-terminus. These findings support a model in which SECISBP2L plays a noncanonical role in regulating gene expression independent of selenoprotein synthesis. Given prior associations between SECISBP2L downregulation or mutation and cancer progression, our data raise the possibility that SECISBP2L modulates cell adhesion and extracellular matrix gene networks relevant to metastatic potential. This work establishes a foundation for further mechanistic studies into SECISBP2L’s role in gene regulation and disease.

## Introduction

Selenocysteine (Sec), often referred to as the 21st amino acid, is found in all domains of life and is encoded by UGA which typically serves as a stop codon. Redefinition of UGA from stop to Sec in eukaryotes requires Sec-tRNA^Sec^, the elongation factor eEFSec, a selenocysteine insertion sequence (SECIS) element in the 3’ UTR, and the SECIS binding protein, SBP2^1^. The SBP2/SECIS complex interacts with the small ribosomal subunit and recruits the eEFSec•GTP•Sec-tRNA^Sec^ ternary complex to the translating ribosome^2^. In humans, 25 selenoproteins result from successful Sec incorporation. Many of these proteins, such as glutathione peroxidases and thioredoxin reductases, are responsible for countering intracellular oxidative stress. Deiodinase enzymes are responsible for proper thyroid hormone metabolism, selenoprotein P (SELENOP) distributes selenium throughout the body, and other selenoproteins have functions that are not fully characterized^3,4^.

Human SBP2 is a 95 kDa protein with an unstructured N-terminus, a central Sec incorporation domain (SID), and a C-terminal L7Ae RNA binding domain (RBD)^5,6^. The N-terminal half of SBP2 is not required for Sec insertion *in vitro* while the C-terminal half containing the SID and RBD is essential. *In vitro* experiments using the rabbit reticulocyte lysate translation system demonstrated that mutations in the RBD that disrupt SECIS binding also impair Sec incorporation^7^. Intriguingly, SID mutations do not impair SECIS binding yet they disrupt Sec insertion^8^. The mutational analysis of SBP2 in *in vitro* systems was essential to dissecting the molecular mechanism that underpins Sec incorporation but did not address its cellular or organismal impacts. The first described human mutations in SBP2 were found in patients presenting with thyroid dysfunction and were also found to have broad decreases in selenoprotein expression^9^. Another report identified patients with SBP2 mutations that exhibit marked reduction in selenoprotein expression, cognitive impairment, growth delays, and increased sensitivity to UV irradiation^10^. A recent report documented aortic aneurysm formation in patients with selenoprotein deficiency caused by SBP2 mutations, and mouse modeling of human SBP2 mutations and tissue-targeted SBP2 knockout have recapitulated partial selenoprotein loss and aortic aneurysm development thus cementing the role of SBP2 in selenoprotein synthesis *in vivo*^11^.

Intriguingly, the human genome encodes an SBP2 paralog called SBP2-like (SECISBP2L). SECISBP2L was discovered by searching a cDNA database and its C-terminal portion, despite having domains homologous to the SBP2 SID and RBD, did not demonstrate Sec incorporation activity *in vitro* ^*6*^. Subsequent research demonstrated that human SECISBP2L could bind to all human SECIS elements, albeit with lower affinity than SBP2, and that full-length SECISBP2L also did not promote Sec incorporation *in vitro*^12^. However, selenoprotein mRNAs co-immunoprecipitated with SECISBP2L from cell lysates suggesting that SECISBP2L may be involved in regulating selenoprotein expression by interacting with their mRNAs. Consistent with mouse data, partial selenoprotein loss was observed upon CRISPR/Cas9-mediated SBP2 knockout in zebrafish embryos. The maintenance of selenoprotein expression upon SBP2 loss could be due to 1) incomplete SBP2 depletion or 2) Sec incorporation directed by SECISBP2L. CRISPR-Cas9 mediated SECISBP2L knockout in zebrafish embryos did not affect selenoprotein expression. However, injection of zebrafish embryos with CRISPR/Cas9-sgRNA complexes targeting both SBP2 and SECISBP2L resulted in complete elimination of detectable selenoprotein synthesis, suggesting that SECISBP2L directs Sec incorporation when SBP2 is impaired^13^. Additionally it was recently demonstrated that full-length, but not C-terminal, SECISBP2L, directs Sec incorporation in the selenoprotein DIO2 in oligodendrocytes^14^. Thus it is possible that SECISBP2L evolved to regulate the expression of specific selenoproteins or regulate the selenoproteome under certain conditions. In the context of human disease, it was recently discovered that the TT genotype at the rs77468143, a SECISBP2L-proximal SNP, is associated with lower SECISBP2L expression and lung adenocarcinoma^15^. Another study found that a Met644Val mutation in SECISBP2L was associated with the spread of clear cell renal carcinoma to lymph nodes^16^. For both of these cancer association studies, the mechanism driving the association remains to be determined.

In this report we sought to further understand the functional differences between SBP2 and SECISBP2L in the human hepatocellular carcinoma cell line (HepG2). This cell line was chosen because it expresses almost the entire selenoproteome, including SELENOP, which has 10 Sec codons. We show for the first time in a human system that targeting SECISBP2L with CRISPR-Cas9 does not affect selenoprotein expression. Surprisingly, bioinformatics analysis indicates that the expression of cell adhesion molecules is affected, introducing a potentially novel role for the N-terminus of SECISBP2L. A caveat of our system is that we detected peptides arising from translation downstream of the CRISPR-targeted site, raising the possibility that gene expression changes are due to the SECISBP2L N-terminus.

## Results

To determine the contributions of SECISBP2L and SBP2 to human cell biology we obtained from Synthego HepG2 cells in which the *SECISBP2* or *SECISBP2L* (hereafter SBP2 and SECISBP2L) loci were targeted using the Cas9/sgRNA system. We independently amplified genomic DNA and sequenced the targeted loci to confirm that SBP2 and SECISBP2L were mutated (Figure 1A). Sanger sequencing of SBP2-targeted cells revealed a single cytosine insertion in exon 4 of SBP2 (Figure 1B). This mutation creates a +1 frameshift premature termination codon and results in loss of SBP2 as observed by western blot and RT-qPCR (Figure 1C). PCR of gDNA from SECISBP2L-targeted cells yielded two bands which upon sequencing revealed an 11 bp deletion in exon 2 and an 813 bp deletion spanning exon 2 and intron 2 (Figure 1B). Both of these mutations are predicted to create frameshifts and premature translation termination. Analysis of mRNA expression by RT-PCR showed that SBP2 mRNA was decreased and SECISBP2L mRNA remained unchanged (Figure 1C). Although we did not observe loss of SECISBP2L mRNA by RT-qPCR, loss of full length SECISBP2L protein was confirmed by western blot, as was SBP2 protein (Figure 1D). The apparent evasion of SECISBP2L mRNA from nonsense-mediated decay suggests that SECISBP2L mRNA may be stabilized by translation initiation downstream of mutation sites. Indeed, quantitative mass spectrometry analysis from these cell lines revealed that C-terminal SECISBP2L peptides are present, so internal translation initiation may be generating an N-terminally truncated version of SECISBP2L (Figure S1).

**Figure 1.**
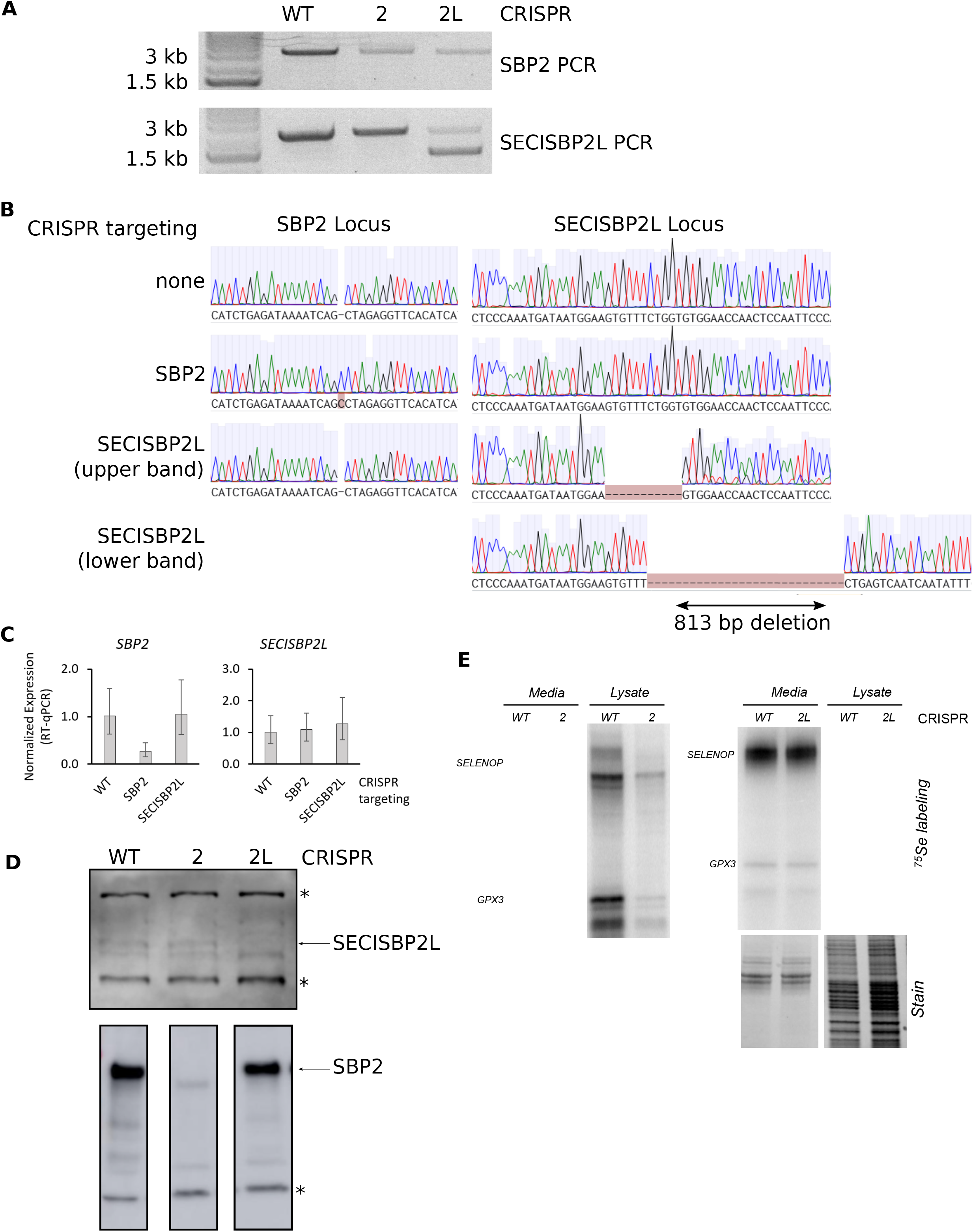
Overview of SBP2 and SECISBP2L CRISPR HepG2 cells. (A). Agarose gel of SBP2 and SECISBP2L PCR products amplified from genomic DNA. (B) Sanger sequencing of the bands shown in panel A and visualized in Benchling. (C) RT-qPCR of SBP2 and SECISBP2L mRNA normalized to GAPDH mRNA. (D) Western blots of SECISBP2L (top) and SBP2 (bottom) from different biological samples. Asterisks indicate non-specific bands to show equal loading. Note that the SBP2 blots were cropped from a larger membrane (see Figure S1). (E) Visualization of selenoproteins by metabolic labeling with ^75^Se.

### SECISBP2L depletion does not alter selenoprotein expression

In order to determine the effect on selenoprotein production, we grew wild-type (WT), SBP2-CRISPR and SECISBP2L-CRISPR cell lines in the presence of ^75^Se to metabolically label selenoproteins. As expected, SBP2-CRISPR cells showed a decrease in overall selenoprotein labeling. In contrast, selenoprotein expression was not affected in SECISBP2L-CRISPR cells (Figure 1D). To gain further insight into the effects of SBP2 or SECISBP2L editing, we conducted RNA sequencing on rRNA-depleted total RNA from cells grown in the presence of supplemental 50 nM sodium selenite. We first assessed expression of selenoproteins. All selenoprotein mRNAs were detected except for those of DIO2, DIO3, GPX6, and SELENOV (Figure 2A). Consistent with the selenium labeling data, SBP2-CRISPR HepG2 cells showed losses of selenoprotein mRNA expression with GPX1, SELENOP, DIO1, and GPX3 being the most strongly affected as determined by differential expression analysis (Figure 2B, Supplemental File 1). This data is generally consistent with the loss of selenoprotein mRNA expression in a mouse liver SBP2 conditional knockout model ^18^(Figure S1). Unsurprisingly, RNAseq selenoprotein mRNA expression in SECISBP2L-CRISPR cells was not strongly affected. However, the mRNAs for DIO1 and SELENOI were downregulated and upregulated ∼30-40%, respectively (Figure 2B, Supplemental File 2). We next purified total RNA from cells cultured in the presence of 50 nM sodium selenite and examined the expression of select selenoprotein mRNAs by RT-qPCR. Consistent with RNAseq, RT-qCPR showed that GPX1, DIO1, and GPX4 mRNAs were downregulated in SBP2-CRISPR cells while SELENOI mRNA was unaffected. In the SECISBP2L-CRISPR cells, RT-qPCR revealed a ∼40% reduction in DIO1 expression while SELENOI, GPX1, and GPX4 mRNAs were unaffected (Figure 2C).

**Figure 2.**
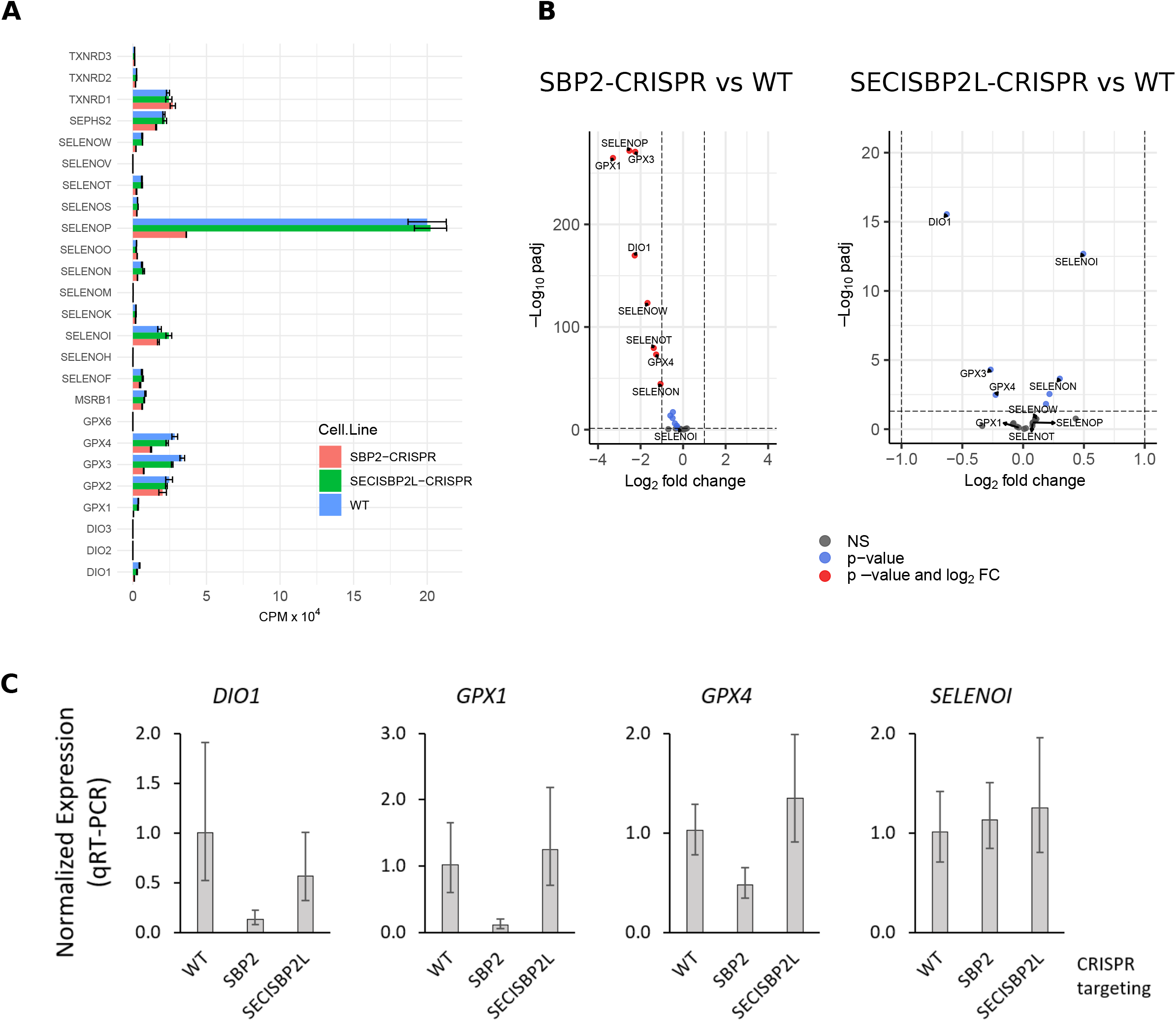
Selenoprotein expression in SBP2 and SECISBP2L CRISPR HepG2 cells. (A) Counts per million mapped reads (CPM) for all selenoprotein genes. Bars are mean ± SD. (B) Volcano plots visualizing DESeq2 output for selenoprotein expression. (C) RT-qPCR analysis of the indicated selenoprotein mRNAs normalized to GAPDH mRNA. Bars are mean ± SD of three replicates.

### SECISBP2L targeting alters expression of ECM and adhesion related genes

As SECISBP2L targeting did not affect selenoprotein transcripts, we sought to further characterize transcriptome changes using over-representation analysis (ORA) of differentially expressed genes. By selecting genes with log2-fold change of >1 and < -1 and p-adjusted of < 0.05 we found that SECISBP2L-depleted cells had a total of 844 differentially expressed genes, comprising 612 upregulated genes and 232 downregulated genes (Figure 3A and Supplemental File 3). Gene ontology (GO) ORA for DEGs revealed GO terms related to cell adhesion and the extra-cellular matrix (Figures 3B and 3C). RT-qPCR analysis of three DEGs, *LTBP1, GPC3*, and *FKBP11*, confirmed their differential expression in SECISBP2L-CRISPR cells (Figure 3D).

**Figure 3.**
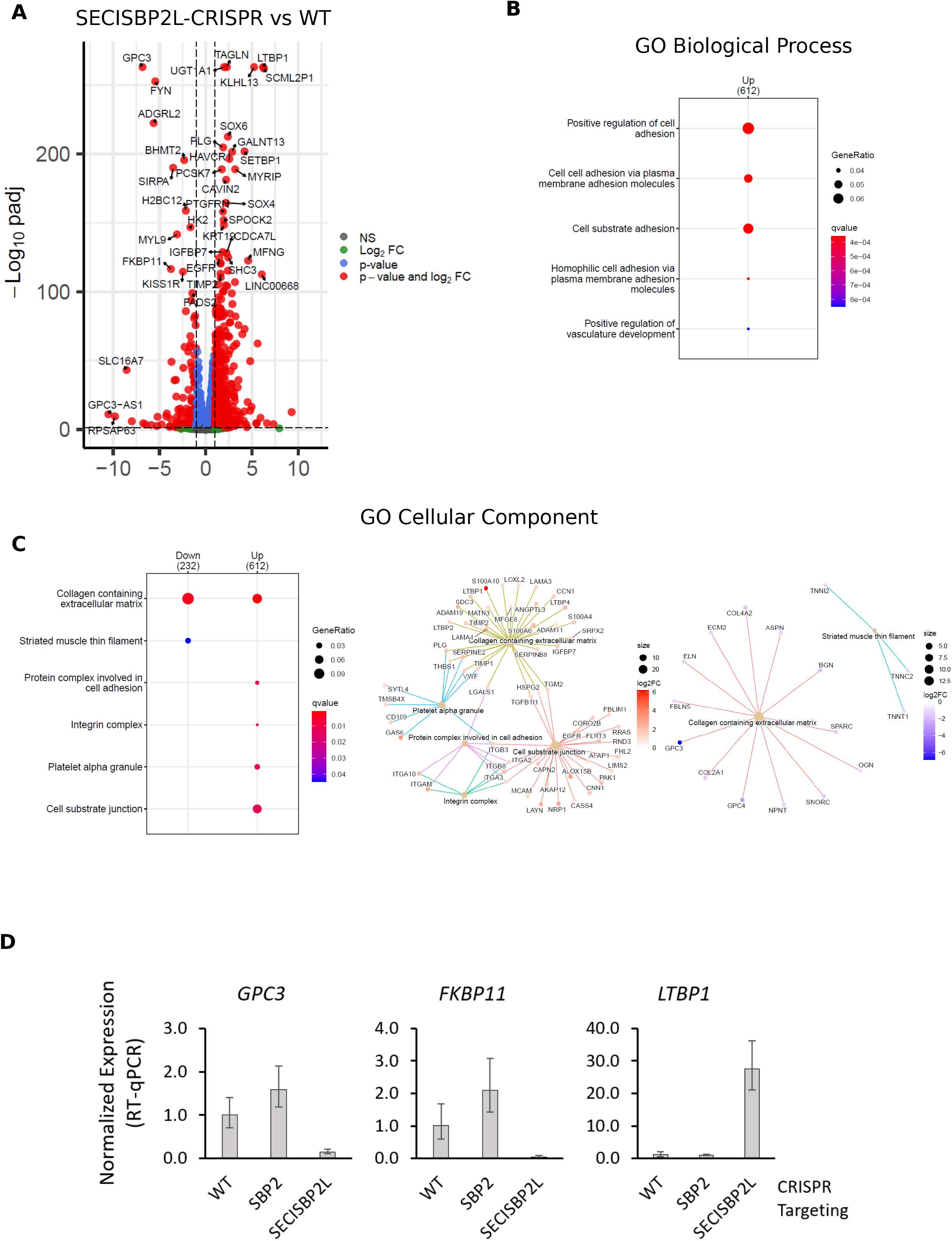
Extracellular Matrix and Adhesion genes are differentially expressed in SECISBP2L-CRISPR HepG2 cells. (A) Volcano plot visualization of differentially expressed genes in SECISBP2L-CRISPR cells. (B) Overrepresentation analysis of DEGs for Gene Ontology Biological Process terms (C) Overrepresentation analysis and gene-concept network plot for Gene Ontology Cellular Component terms. (D) RT-qPCR analysis for the indicated mRNAs normalized to GAPDH. Bars are mean ± SD of three replicates.

### SBP2 depletion results results in a transcriptome distinct from that of SECISBP2L depletion

Analyzing DEGs in SBP2-depleted cells using the same fold-change and statistical cutoffs revealed a markedly smaller set of DEGs with 116 upregulated genes and 195 downregulated genes (Figure 4A, Supplemental File 1). GO Biological Process ORA revealed enriched terms relating to steroid metabolism, fatty acid metabolism, and anion transport among downregulated genes (Figures 4B and 4C). Additionally, the GO Biological Process terms for zinc and copper responses were enriched in upregulated genes (Figures 4B and 4C). No GO Cellular Component terms were enriched in the upregulated DEGs from SBP2-depleted samples. The GO Cellular Component term ‘collagen containing extracellular matrix’ was enriched among the downregulated genes, albeit with low statistical significance. Consistent with the divergences observed by ORA, there are only small overlaps among the DEGs in SECISBP2L-depleted and SBP2-depleted cells. Forty genes are upregulated in both SBP2 and SECISBP2L targeted cells and thirty-six downregulated genes are common between both mutant cell lines. Thirteen genes upregulated in the SBP2-CRISPR cells were also downregulated in the SECISBP2L-CRISPR cells. Forty-five genes downregulated in SBP2-CRISPR cells were upregulated in the SECISBP2L-CRISPR line (Figure 4D).

**Figure 4.**
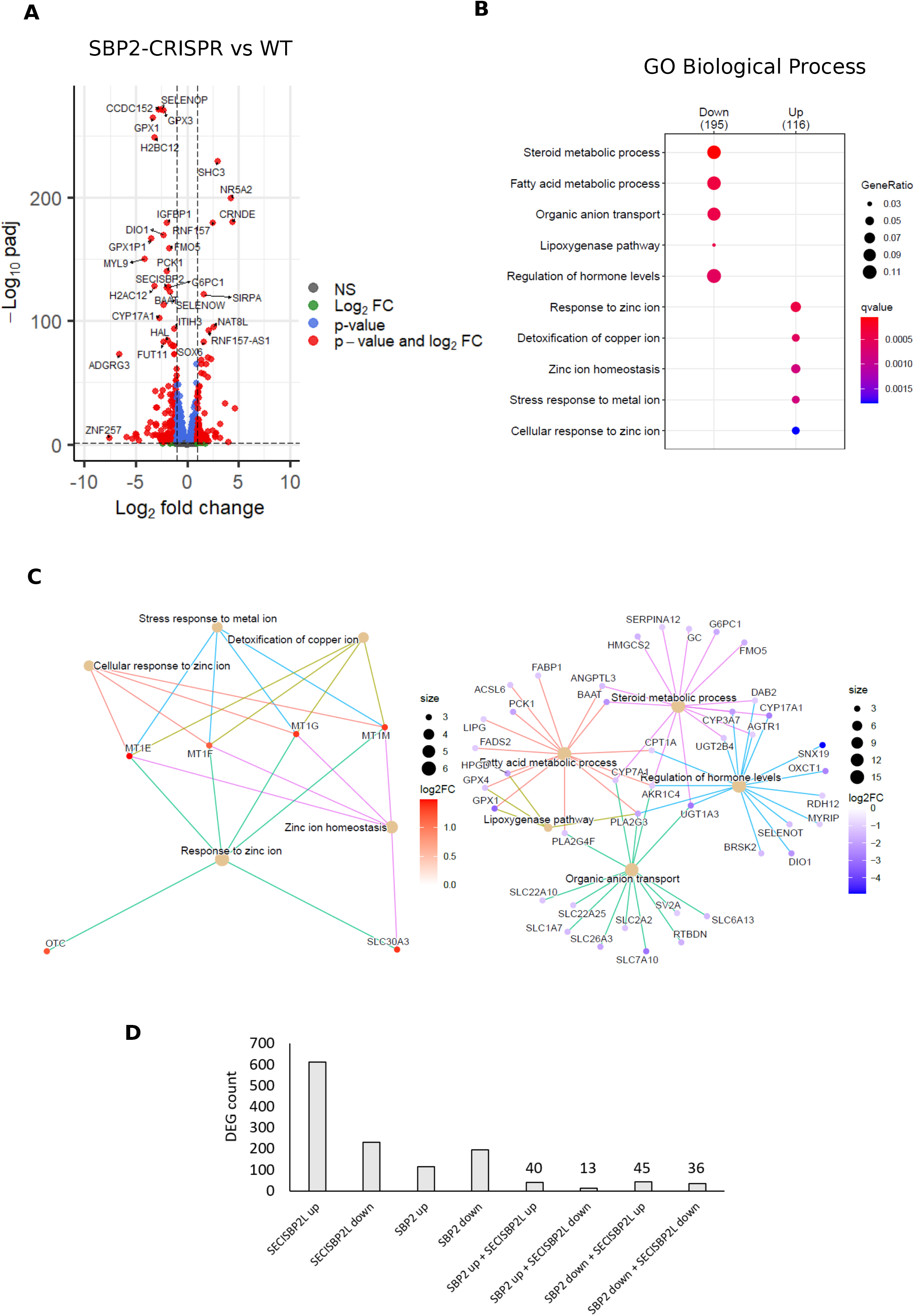
Divergent transcriptome signature in SBP2-CRISPR and SBP2L-CRISPR cells. (A) Volcano plot visualization of differentially expressed genes in SBP2-CRISPR cells. (B) Overrepresentation analysis for GO Biological Process terms among DEGs from SBP2-CRISPR cells. (C) Gene-concept network plot for the GO terms in panel B. (D) DEG counts and overlap between SBP2-CRISPR and SECISBP2L-CRISPR HepG2 cells.

## Discussion

Here we presented the first transcriptomic comparison resulting from CRISPR editing of the paralogous proteins SBP2 and SECISBP2L, which resulted in complete depletion of SBP2 and truncation of the N-termninus for SECISBP2L. The expected loss of selenoprotein transcripts in SBP2-CRISPR cells and minimal effect on selenoprotein transcripts in SECISBP2L-CRISPR cells was consistent with existing literature. However, recent results in a zebrafish model suggest that SECISBP2L and SBP2 co-depletion abrogates selenoprotein expression whereas single SBP knockout had minimal effects. Thus in our human HepG2 model, the possibility remains that incomplete loss of selenoprotein expression in SBP2-CRISPR cells is due to activation of SECISBP2L. The large DEG set discovered upon loss of SECISBP2L expression also raises interesting questions. Which of these genes are the result of transcriptional changes and which gene expression changes are post-transcriptional and controlled by the RNA binding activity of SECISBP2L? Owing to its L7Ae RNA binding motif it is reasonable to speculate that SECISBP2L binds to some mRNAs uncovered in our bioinformatic analysis via a SECIS-like structure. The enrichments of ECM and adhesion related genes among the up- and downregulated DEGs that arise from the loss of SECISBP2L are particularly interesting in light of the recent reports connecting lower SECISBP2L expression with lung cancer and an SBP2L mutation with metastasis of clear cell renal carcinoma^15,16^. It is tempting to speculate the downregulation or mutation of SECISBP2L in cancer enhances metastasis through changes in the expression of ECM and adhesion gene clusters. A detailed mechanistic understanding of these phenomena and their implications for basic biology and disease awaits further experimentation.

## Materials and Methods

### Cells and Cell Culture

Wild-type HepG2 cells were obtained from ATCC. CRISPR-targeted SBP2 and SECISBP2L HepG2 lines were purchased from Synthego (Redwood City, CA, USA). Cells were maintained in EMEM with 10% FBS and 50 nM sodium selenite.

### SBP2 and SECISBP2L genotyping

Genomic DNA (gDNA) was purified using QuickExtract DNA Extraction Solution (Biosearch). gDNA regions of interest were amplified using Accuprime HiFi Taq and the following primers: [SBP2] 5’-TCCTTATACCCTTGACTCCACA-3’ and 5’-TCCTGTCTGTTCGCTTATGG-3’; [SECISBP2L] 5’-TGTCAGCTGAGGTGGAGCCATT-3’ and 5’-TACTCTGTAGAAACAGGCGGCTGAGCAG-3’. PCR reactions were resolved by 1% agarose gel electrophoresis and bands were excised and purified with the Purelink Gel Extraction Kit (Invitrogen). Purified PCR products were analyzed by Sanger sequencing using 5’-TCCTGTCTGTTCGCTTATGG-3’ and 5’-TGTCAGCTGAGGTGGAGCCATT-3’ primers for SBP2 and SECISBP2L, respectively.

### RNA sequencing and data processing

Cells, in triplicate, were lysed in 20 mM HEPES pH 7.5, 150 mM NaCl, 1X c*0*mplete EDTA-free protease inhibitor cocktail (Roche), 1% NP-40 and incubated on ice for 20 min. Lysates were then cleared by centrifugation for 15 min at 14,000 x g, 4 °C. RNA for RNA sequencing was purified from the clarified lysate using Trizol LS (Life Technologies). Libraries were prepared by the Rutgers University Genomics core facility and 150 bp pair-end sequencing was run on an Illumina HiSeq. One sample each from SBP2-CRISPR and SECISBP2L-CRISPR cells has library sizes that were noticeably shorter by analysis on an Agilent Tapestation and were therefore omitted from data analysis. Illumina adapter sequences were removed using TrimGalore^19^ and mapped to the human genome (Hg38) using STAR^20^. Gene expression was estimated using featureCounts^21^. Differential gene expression was called using DESeq2^22^ with the following cutoffs DEG calling: log2foldchange = 1/-1, Benjamini-Hochberg FDR < 0.05, baseMean expression > 10 . DESeq2 analysis, ORA, and gene-concept network plots were conducted within the RNASeqChef web app^23^. DESeq2 output from the RNAseq Chef server required that log2foldchange to be corrected by multiplying the results by -1. For analysis of published RNASeq data, FASTQ files from GSE84122 were imported to Galaxy and processed as described above.

### ^75^Se Metabolic Labeling

WT or CRISPR edited HepG2 cells were grown in 96-well plates to approximately 70% confluence when the medium was changed to serum-free medium supplemented with 100 nM ^75^Se (University of Missouri Research Reactor; specific activity of ∼500 Ci/g) or 50 uCi/ml 35S Met/Cys (Perkin Elmer). After 24 hours of labeling, cells were lysed in 100ul of 2X SDS sample buffer (125 mM Tris-HCl pH 6.8, 4% (w/v) SDS, 0.02% (w/v) bromophenol blue and 20% (v/v) glycerol) and heated at 95 °C for 5 minutes. Proteins were resolved by 15% SDS gels. The gels were dried and exposed to a phosphorimage screen for 24 hrs before being developed on the Typhoon FLA7000 (Cytiva).

### Western Blotting

Cells were lysed in in RIPA buffer (50 mM Tris-HCl pH 7.5, 150 mM NaCl, 1% NP-40, 0.1% SDS, and 0.1% deoxycholate) supplemented with 1X complete EDTA-free protease inhibitor cocktail. Lysates were cleared by centrifugation at 20,000 x g for 20 min at 4 °C. Clarified supernatants were transferred to new tubes and protein concentrations determined using 660nm Protein Assay reagent (Pierce) and BSA standard. Samples for SBP2 and SECISBP2L blotting were resolved by 8% SDS-PAGE and transferred to nitrocellulose. SECISBP2L was detected using chicken anti-SECISBP2L (Invitrogen; 1:500) and HRP-conjugated goat anti-chicken (1:20,000). SBP2 was detected using rabbit anti-SBP2 (Proteintech) and HRP-conjugated goat anti-rabbit. All blots were detected using SuperSignal West Femto (Pierce).

### Mass spectrometry

Cells, in triplicate, were lysed in RIPA buffer as described above and quantitative mass spectrometry was performed on a Bruker TIMS/TOF HT mass spectrometer by the Rutgers Biological Mass Spectrometry facility.

### RT-qPCR

Total RNA was purified from cells using Trizol according the manufacturer’s protocol and resuspended in water. RNA samples were then digested with DNase I for 15 min at 37 °C and re-extracted with 1 volume of phenol:chloroform:isoamyl alcohol (25:24:1) and precipitated with ethanol. RNA pellets were resuspended in water, quantified by Nanodrop, and equal masses of RNA were reverse-transcribed with SuperScript IV reverse transcriptase primed with random hexamers according to the manufacturer’s protocols. cDNAs were diluted with water and used as templates for qPCR with Power SYBR green mastermix (Applied Biosystems). qPCR primers are listed in Supplemental File 4). Each of three biological samples were analyzed with three technical replicates and data were normalized to GAPDH using the delta-delta Ct method.

