## Supplementary figures and images for "RNA Sequencing Reveals Divergent Transcriptome Changes in SBP2 and SECISBP2L Depleted Cell Lines"

### Figure S1

#### Figure S1

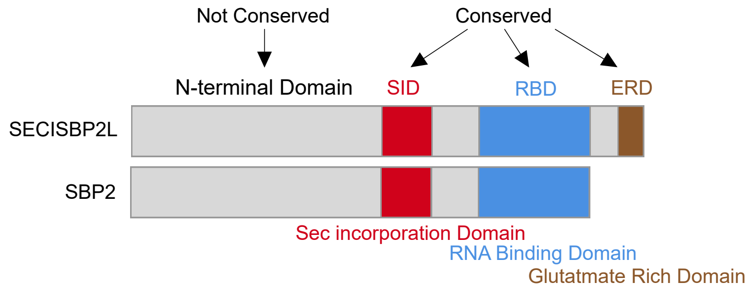


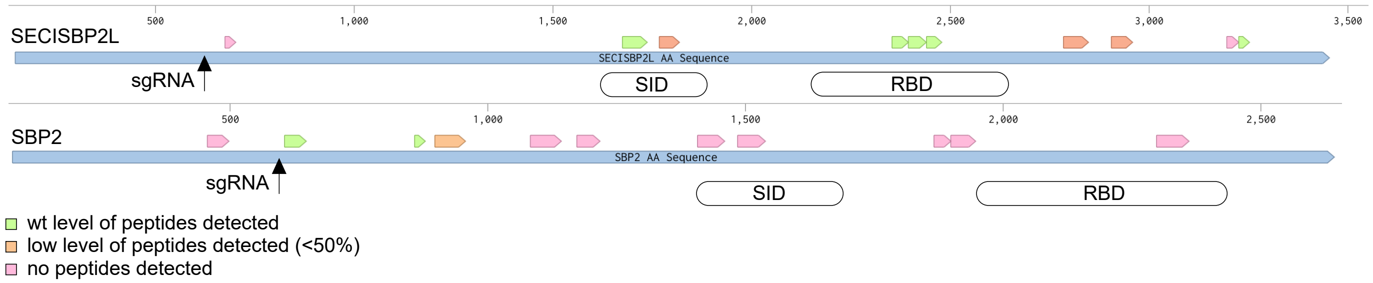


SBP2 western blot

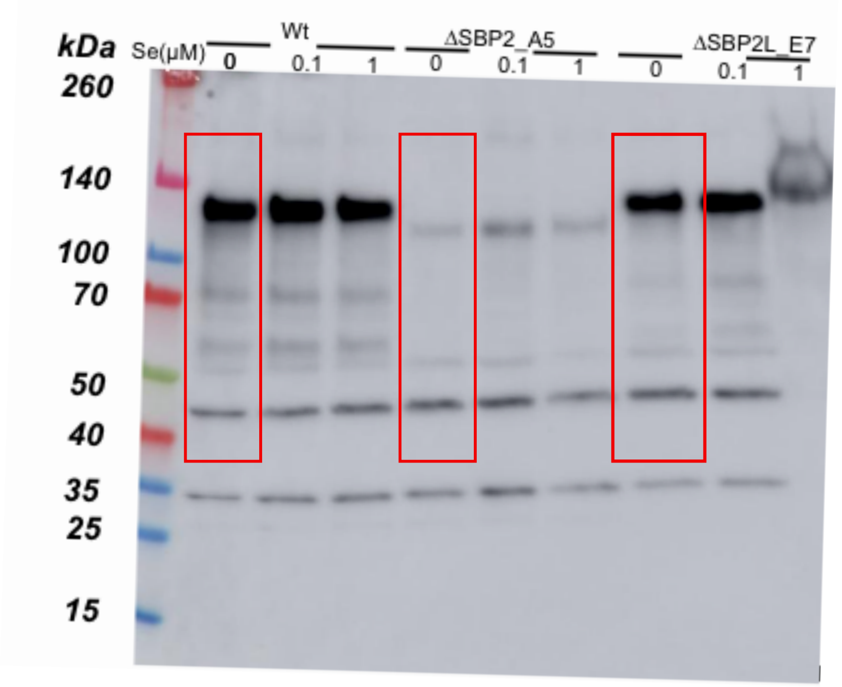

### Figure S2

Figure S2

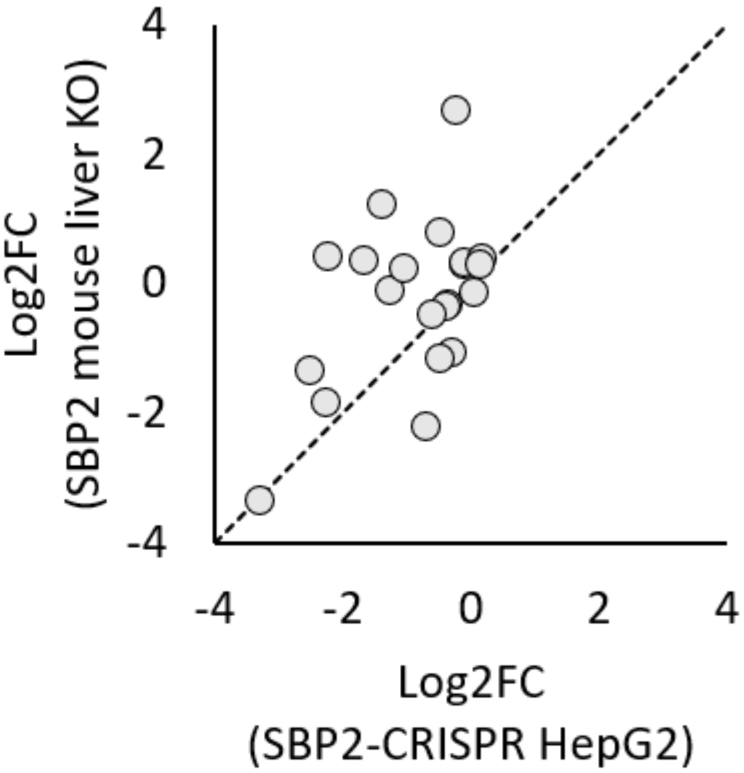


Pearson correlation = 0.48
